# Improved stability of an engineered function using adapted bacterial strains

**DOI:** 10.1101/2020.03.05.979385

**Authors:** Drew S. Tack, Peter D. Tonner, Elena Musteata, Vanya Paralanov, David Ross

**Author notes:** Corresponding Author, E. mail, 100 Bureau Dr. Gaithersburg, Maryland, 20899, USA, Peter D. Tonner, Elena Musteata, Vanya Paralanov.

## Abstract

Engineering useful functions into cells is one of the primary goals of synthetic biology. However, engineering novel functions that remain stable for multiple generations remains a significant challenge. Here we report the importance of host fitness on the stability of an engineered function. We find that the initial fitness of the host cell affects the stability of the engineered function. We demonstrate that adapting a strain to the intended growth condition increases fitness and in turn improves the stability of the engineered function over hundreds of generations. This approach offers a simple and effective method to increase the stability of engineered functions without genomic modification or additional engineering and will be useful in improving the stability of novel, engineered functions in living cells.

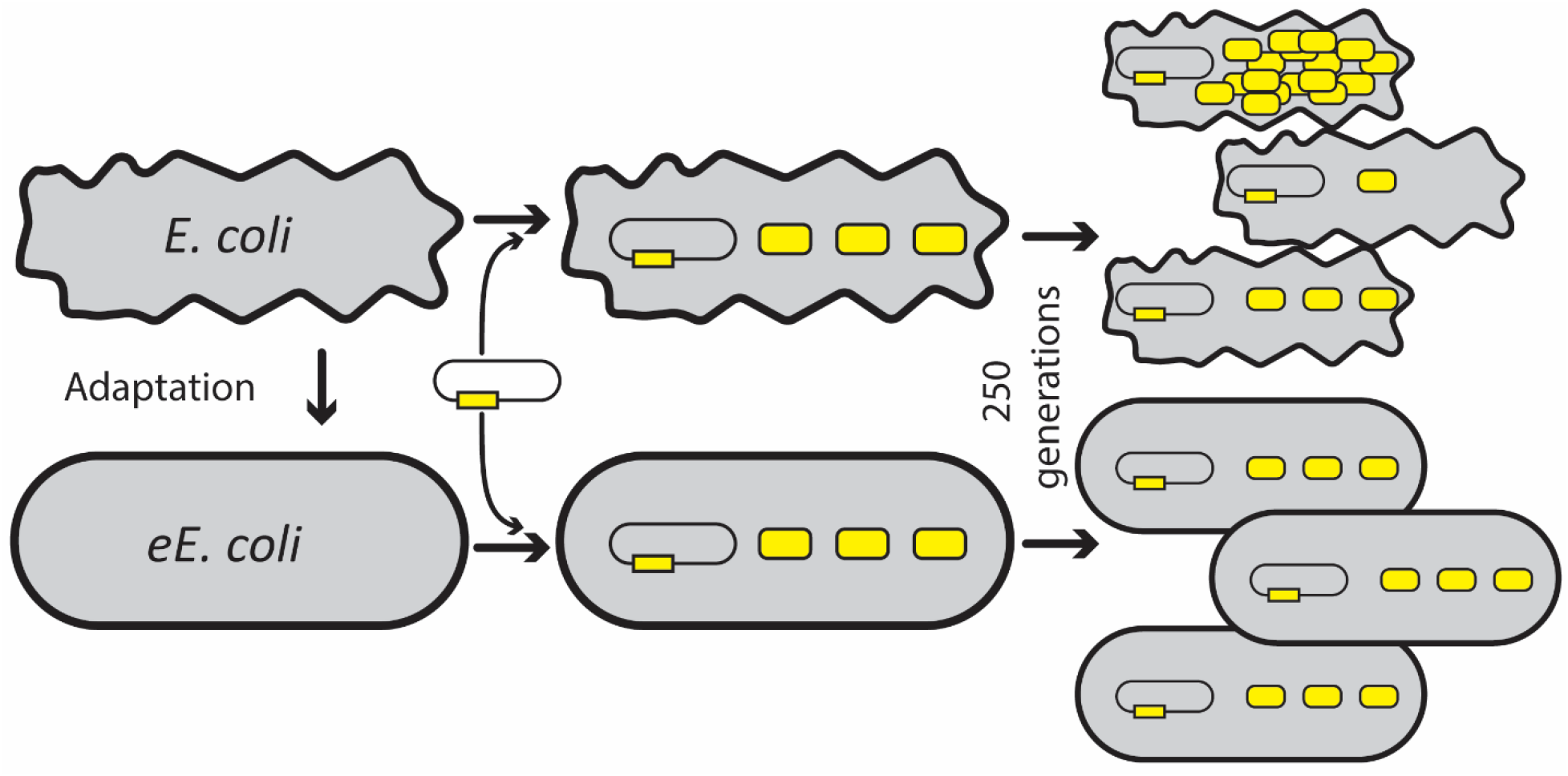

## Introduction

Advances in synthetic biology over the last few decades have enabled the engineering of cells to perform a wide range of new and useful functions. However, engineered functions are often unstable, with variability of the function over generations. The stability of engineered functions has been improved by regulating genetic elements^1,2^, reducing mutation rates of the host organism,^3-6^ entangling essential and nonessential genetic features^7^, and integrating genetic components into the host genome^8,9^. These approaches are engineering efforts themselves and require significant investments in time and resources. Simple, accessible, and universal approaches to increase the stability of engineered functions will greatly facilitate the application of synthetic biology to problems where functional stability is important.

Here, we investigate the stability of an engineered function over hundreds of generations in several strains of *Escherichia coli*, a workhorse model organism for many research, industrial, and medical applications. We identify the fitness of the host strain as an important determinant in maintaining the engineered function. We demonstrate that adapting a strain to the intended growth condition prior to encoding the engineered function improves strain fitness and increases the stability of the engineered function. This approach offers a new and complementary tool for engineering cellular functions for industrial and environmental applications that require functional stability over evolutionary timescales.

## Results and Discussion

We selected five *E. coli* strains to explore the stability of engineered functions in different hosts, including the cultivated wild-type MG1655, a nontoxigenic serotype O157:H7, and three commercially available cloning strains: TOP10, DH5α, and NEB Turbo. We used fluorescence as an engineered function that could be measured rapidly and accurately with flow cytometry at the single cell level. To encode fluorescence, we transformed each strain with plasmid pRPU (Supplementary Figure 1a), which encodes a constitutively expressed Yellow Fluorescent Protein (YFP) and provides a reference for relative gene expression measurements^10^. To investigate the stability of each strain, we passaged each containing pRPU at a rate of 10 generations per day for 25 days, *i.e.* 1000-fold dilutions at 24-hour intervals for 250 generations, providing an evolutionary timeframe for loss of function to occur^11^. We measured the YFP expression of the populations during exponential growth using flow cytometry (Figure 1, left panels). The results show that the stability of YFP expression over 250 generations was strain dependent (supplementary table 1), with some strains exhibiting consistent expression of YFP throughout the course of passaging (*e.g.* TOP10) and others exhibiting high variability over generations and between replicates (*e.g.* DH5α). The observed differences in stability of the engineered function between strains suggests the choice of host is an important factor in the stability of engineered functions and must be considered for the desired application.

**Figure 1.**
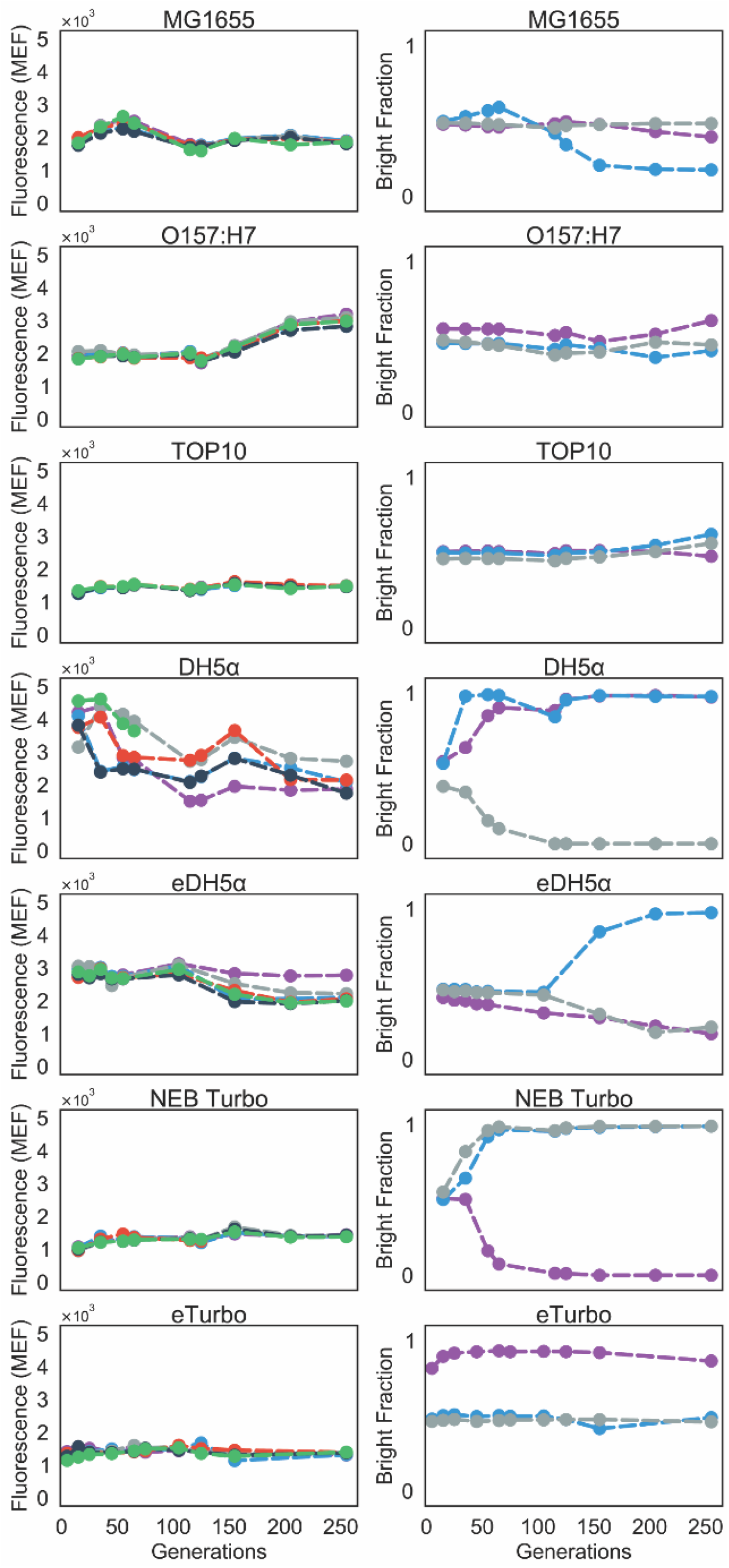
Stability of YFP expression in seven different *E. coli* strains. Left, stability assays measured the fluorescence of populations constitutively expressing YFP over 250 generations (mode, reported as molecules of equivalent fluorophore, MEF). Some strains maintained a consistent expression level (e.g. TOP10), and others had high variability of YFP expression over generations and between biological replicates (e.g. DH5α). Right, competitive fitness assays between cells expressing YFP (‘bright’) and cells not expressing YFP (‘dark’). Populations began as equal mixtures of bright and dark cells. Stable strains remained as mixed populations over 250 generations (e.g. TOP10), while unstable strains were driven to populations dominated by either bright or dark cells (e.g. DH5α). Both assays were completed in biological triplicate. Stability assays also display fluorescence intensity of the bright populations from the competitive fitness assays. Error bars represent standard deviation and are within markers.

The stability of engineered functions has previously been correlated with the fitness cost of the function^12,13^. To estimate the fitness cost of YFP expression, we created a negative control, plasmid pRPU-neg (Supplementary Figure 1b), by inactivating the promoter and ribosome binding site of YFP on plasmid pRPU. We conducted competitive fitness assays by mixing populations of cells containing pRPU (‘bright’) and cells containing pRPU-neg (‘dark’) at a 1:1 ratio. We then passaged the mixed cultures as described above and measured the ratio of bright and dark cells over time with flow cytometry. Strains MG1655, TOP10, and O157:H7 remained close to a 1:1 ratio throughout 250 generations (Figure 1, right panels). In contrast, competitive fitness assays of NEB Turbo and DH5α quickly resulted in populations dominated either by dark or by bright cells. Since competitive fitness assays were stable in three of the strains, and four of the six replicates of unstable strains were driven to bright populations, we conclude that YFP expression has a minimal impact on fitness, and that deactivating the function does not greatly improve fitness. Therefore, other factors likely impact the outcomes of the stability and competitive fitness assays.

**Figure 2.**
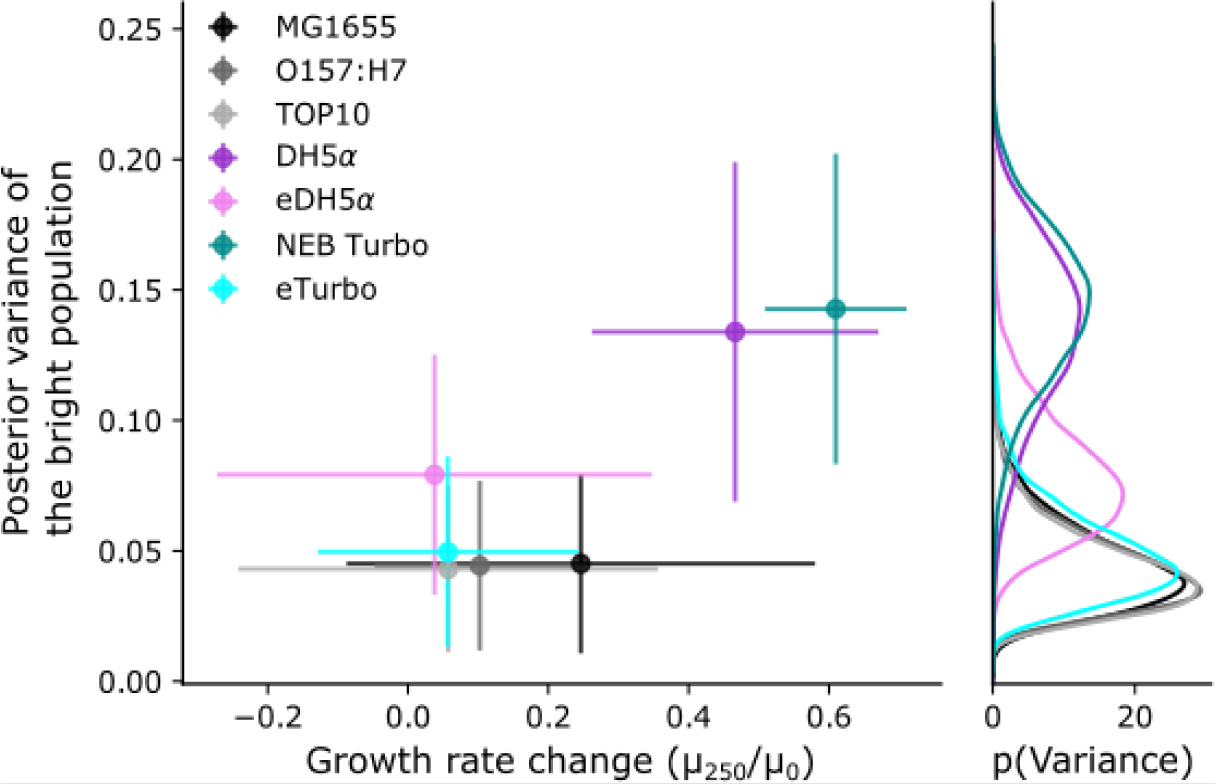
Growth rate changes and stability of seven *E. coli* strains analyzed with a hierarchical Bayesian inference model. Left panel, the posterior variance of the bright population from the competitive fitness assays is plotted against the growth rate change for each strain. High posterior variance corresponds to instability and unpredictable outcomes while low variance corresponds to higher stability with predictable outcomes. Right panel, the distribution of the posterior variance from the hierarchical model of stability, corresponding to the vertical error bars of left plot. Strains which had larger changes in growth rate had higher posterior variance in the outcomes of the competitive fitness assay, indicating lower stability and

To determine what other factors are important for the stability of the engineered function, we measured growth rates as an indicator of fitness at the beginning (generation 0, μ_0_) and conclusion (generation 250, μ_250_) of both the stability and competitive fitness assays (Supplementary Table 2). The two *E. coli* strains that were unstable (DH5α and NEB Turbo) had significantly increased growth rates during the competitive fitness assays (>40%, P<.001 for both comparisons, Figure 2, Supplementary Table 2, Supplementary Figures 2 and 3). In contrast, the three *E. coli* strains that were stable during competitive fitness assays (MG1655, TOP10, and *E.c.*O157:H7) exhibited modest increases in growth rates (0-20%, Figure 2).

Based on these results, we hypothesized that the initial strain fitness was the most important determinant of stability in our assays, and that adapting slow-growing cells to the growth condition would increase fitness and, in turn, the stability of the engineered function. To test this hypothesis, we adapted DH5α and NEB Turbo before introducing the plasmid pRPU, by growing each in the enriched M9 media used for stability and competitive fitness assays. To adapt the DH5α and NEB Turbo strains, we grew each strain for 100 generations in the enriched M9 media (see methods). Clonal isolates from the conclusion of the adaptation, eDH5α and eTurbo, had significantly increased growths rates in enriched M9 media compared to ancestral strains, (P<0.0001 for both comparisons, Supplementary Tables 3 and 4, Supplementary Figure 4). In contrast, the fitness of both adapted strains in LB media was not different than ancestral strains (Supplementary Tables 3 and 4, Supplementary Figure 4). We assayed stability and competitive fitness with these adapted strains and found increased stability of YFP expression when compared to their ancestral strains (Figure 1, left panels, Figure 2, Supplementary Table 2, compare DH5α to eDH5α and NEB Turbo to eTurbo). These results support our hypothesis that the stability of the engineered function is affected by fitness, and that adapting the strains prior to introducing the engineered function improves the stability of that function.

The adapted strains also showed increased stability in the competitive fitness assays, with five of the six replicates from the adapted strains remaining as mixed populations of bright and dark cells (Figure 1, compare competitive fitness assays of DH5α to eDH5α and NEB Turbo to eTurbo). To provide a quantitative measure of stability, we considered the predictability of the competitive fitness assays and modeled the outcomes as a hierarchical process with Bayesian inference^14^. We analyzed strain stability by considering the variance of the final bright fraction of the population. High variance corresponds to unpredictable fixation of the population to either fully dark or fully bright, while low variance corresponds to a predictable outcome with a stable, mixed population. Our analysis confirms the previous observation of high stability in strains MG1655, O157:H7, and TOP10, and lower stability in strains DH5α and NEB Turbo (Figure 2). This analysis also found the adapted strains eDH5α and eTurbo had improved stability compared to ancestral strains (Figure 2). In summary, adapted strains had smaller changes in their growth rates than their ancestral strains and, in turn, were more stable during the competitive fitness assays.

In this study, we investigated the stability of an engineered function in seven strains of *E. coli.* While encoding the function did not measurably affect host fitness, the engineered function was more stable in strains with higher fitness. In addition, adapting strains to the intended growth conditions rapidly increased their fitness and improved the stability of the engineered function when the function is introduced after adaptation. Reduced variability in the function over hundreds of generations as well as increased predictability in competitive fitness assays demonstrate improved stability using adapted strains. These results indicate that a brief adaptation to the intended growth condition can rapidly improve the fitness of a desired host strain and can greatly improve the stability of engineered functions. The approach described here is simple and effective and requires no prior knowledge or additional engineering. Additionally, this approach should be generalizable to any microorganism or growth condition and can be implemented in industrial and other applications.

## Methods

### Growth Conditions

Enriched M9 media was prepared using M9 salts (Difco M9 Minimal Salts) supplemented with 2 mmol L^−1^ magnesium sulfate (Sigma), 100 μmol L^−1^ calcium chloride (Sigma), 0.4% glucose (sigma), and 0.2% Omnipur Casamino Acid (Calbiochem). When required for plasmid maintenance, M9 media was supplemented with kanamycin (Gibco) at 50 μg mL^−1^. LB Miller media was prepared as directed (Fisher BioReagents). Bacto Agar (BD Biosciences) was added to 1.5% to produce solid media.

*E. coli* strains were grown in vertical 14 mL culture tubes (Falcon), shaking at 300 rpm and maintained at 37°C.

### Strains and Plasmids

Wild-type *E. coli* strains MG1655 (ATCC 47076) and O157:H7 (ATCC 700728) were acquired from the American Type Culture Collection (ATCC). One Shot TOP10 *E. coli* was acquired from Invitrogen (C404052). *E. coli* strains DH5α (NEB C2987I) and NEB Turbo (NEB C2984I) were acquired from New England Biolabs.

All plasmid transformations were accomplished with electroporation. Strains were made electrocompetent by growth in LB media at 37°C to mid-exponential phase (OD~0.5). Cultures were then chilled on ice for 15 minutes, followed by centrifugation and serial washing using chilled 10% glycerol. Cells were transformed with appropriate plasmid in 1 mm Bio-Rad Gene Pulser Cuvettes at 1.8 kV using Eporator (Eppendorf). Transformed cells were recovered in LB media for 1 hour at 37°C and plated on LB-agar supplemented with kanamycin.

Plasmid pRPU was a gift from the laboratory of Professor Christopher Voigt. Plasmid pRPU-neg was constructed using primers DT.1 (5’-CCGGTCTGATGAGTCCGTGAGGACGAAACA-GCCTCTACAAATAATTTTGTTTAATACTTGTGTTTGTGGGGTTTTACTAGATGGTGA-GCAAGGGCGAGG-3’) and DT.2 (5’-CTGTTTCGTCCTCACGGACTCATCAGACCGGAA-AGCACATCCGGTGACAGCTTCAGGCTAGCCACCACCCTAGGACTGAGCTAGCTGTA-AAAGTTAGGG-3’) to disrupt the −10 region of the promoter and RBS sequence of the YFP coding DNA sequence on pRPU. PCR product was circularized using the Gibson assembly and sequences were verified using Sanger sequencing (Psomagen).

Adapted strains eDH5α and eTurbo were generated from DH5α and NEB Turbo, respectively. Single clones of *E. coli* strains NEB Turbo and DH5α were isolated on LB-agar and grown overnight in LB media. The same overnight cultures were passaged by transferring 0.25 μL of culture into 40 mL of enriched M9 media. Cultures were incubated at 37°C while shaking until full density was reached (36 hours) and passaged again by diluting 0.25 μL of culture into 40 mL of enriched M9 media. This process was completed a total of six times, allowing for 100 doublings of the original inoculum in enriched M9 media. At the conclusion, an aliquot of each culture was plated on enriched M9 media and colonies were selected for characterization and growth curve measurements (Supplementary Figure 4).

### Passaging

For each strain used, three colonies containing plasmid pRPU and three colonies containing plasmid pRPU-neg were selected after transformation and used to inoculate liquid cultures. Replicates were grown in LB media for 16 hours, after which an aliquot from each was used to make glycerol stocks and stored at −80°C. Overnight cultures were also used to inoculate enriched M9 media for generation 0 of the stability and competitive fitness assays.

During stability and competitive fitness assays, cultures were passaged at 24-hour intervals by transferring 3 μL of the previous day’s culture into 3 mL of new enriched M9 media supplemented with kanamycin. Glycerol stocks of cultures were stored at generation 0, generation 70, generation 150, and generation 250.

### Flow Cytometry

On measurements days, cultures were passaged to new enriched M9 media as described above. 3.5 hours after dilution, cultures were removed from the incubator and a sample of each was removed for flow cytometry measurements. Cultures were immediately returned to shaking incubator. Samples removed from cultures were diluted into phosphate buffered saline (PBS pH 7.4, Invitrogen) supplemented with chloramphenicol (170 μg mL^−1^, Sigma) to halt protein translation. Samples were then stored at room temperature for 1 hour to allow YFP maturation. Culture dilutions were adjusted to measure approximately 10^5^ cytometry events per sample.

The resulting diluted samples were measured on an Attune NxT flow cytometer with 488 nm excitation laser and a 530 nm ± 15 nm bandpass emission filter. Blank samples were measured with each batch of cell measurements, and an automated gating algorithm discriminated cell events from non-cell events. A second gating algorithm identified singlet cell events and excluded multiplet cell events. All subsequent analysis was performed using the singlet cell event data (Supplementary Figure 5). Fluorescence data was calibrated to molecules of equivalent fluorophore (MEF) using Spherotech Rainbow Calibration Beads (lot AJ01)^15^.

The flow cytometry measurement of generation 115 was delayed 3 hours (stored for a total of 4 hours in PBS with 170 μg mL^−1^ chloramphenicol), while troubleshooting an autosampler error.

### Growth Rates

Growth rates were determined after the conclusion of the stability assays and competitive fitness assays by reviving stored samples from generation 0 and generation 250 of each assay. Frozen glycerol stocks were used to inoculate initial cultures in enriched M9 media in 96 well plates (500 μL culture volume per well in clear, square-well assay plates, Brooks Life Sciences 4ti-0255). Cultures were grown overnight at 37°C shaking at 1300 rpm. Overnight cultures were used to inoculate triplicate plates (A, B, C) by transferring 1 μL of overnight culture into 500 μL of media in 96 well plates. Growth curves were measured by monitoring OD_600_ every three minutes for 12 hours using a Tecan Spark 20M running Spark Control V2.1, paired with large humidity cassette. Growth rates were calculated using early exponential growth (OD_600_ measurements between .05 and .1 above background), fitting to the longest contiguous region of data within these values.

### Hierarchical stability model

For the hierarchical model, we modeled the populations of bright cells (cells containing pRPU) and the population of dark cells (cells containing pRPU-neg) in each experiment with a beta distribution:

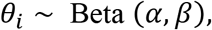

where *θ_i_* is the fraction of bright cells, and the hyperparameters *α* and *β* were given Gamma hyperpriors with the shape and rate parameters 2 and 1, respectively. The bright fraction parameter was then used to model the number of fluorescent cells in each assay as a binomial distribution. The posterior variance of the fraction of bright population (variance *θ_i_*) was used as an estimate of strain stability. Inference was performed using Hamiltonian Monte-Carlo, as provided by Stan^14^.

## Supporting information

Five supplemental figures and five supplemental tables

## Author Contribution

DST designed and performed experiments, analyzed data, and wrote the manuscript. PDT performed hierarchical Bayesian modeling and wrote the manuscript. EM performed experiments and analyzed data. VP designed and performed experiments and wrote the manuscript. DR designed and performed experiments, analyzed data, and wrote the manuscript.

## Disclaimer

Certain commercial equipment, instruments, or materials are identified in this paper in order to specify the experimental procedure adequately. Such identification is not intended to imply recommendation or endorsement by the National Institute of Standards and Technology, nor is it intended to imply that the materials or equipment identified are necessarily the best available for the purpose.

## Acknowledgements

We acknowledge financial support from the NIST SRM Service Development program. This work was supported by a National Research Council Post-Doctoral Research Associateship to DST.

