## Supplementary material for "Improved stability of an engineered function using adapted bacterial strains": Five supplemental figures and five supplemental tables

**Supplementary Figure 1**: Maps of plasmids used in this study. (a) Plasmid pRPU encoded fluorescence function into *E. coli*. Plasmid pRPU contains the YFP coding DNA sequence (CDS) under control of the constitutive promoter $P_{J23101}$. The mRNA transcript from $P_{J23101}$ is insulated with the riboJ insulator and terminated downstream of the YFP CDS with the transcriptional terminator $T_{L3S2P21}$. The plasmid contains the p15A origin of replication and the kanamycin resistance marker. (b) Plasmid pRPU-neg, used for competitive fitness assays, was constructed by inactivated the $P_{J23101}$ promoter and corresponding RBS.

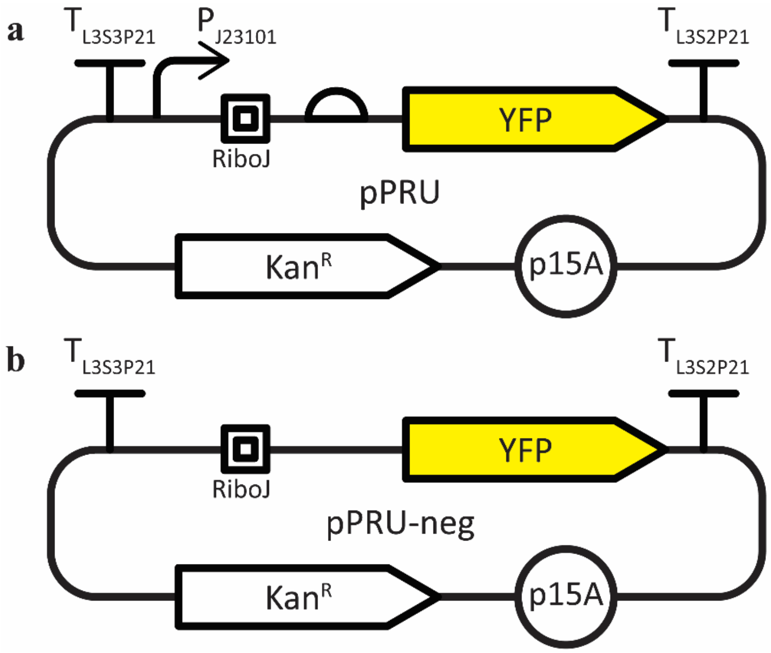

**Supplementary Figure 2**: Growth curves of *E. coli* strains at generation 0 and generation 250 of stability assays. *E. coli* strains containing pRPU and grown in M9 media with kanamycin. Growth curves were measured from samples of each culture at generation 0 (gray curves) and generation 250 (black curves). Each biological replicate (A, B, C) was measured in technical triplicate. Some curves overlap.

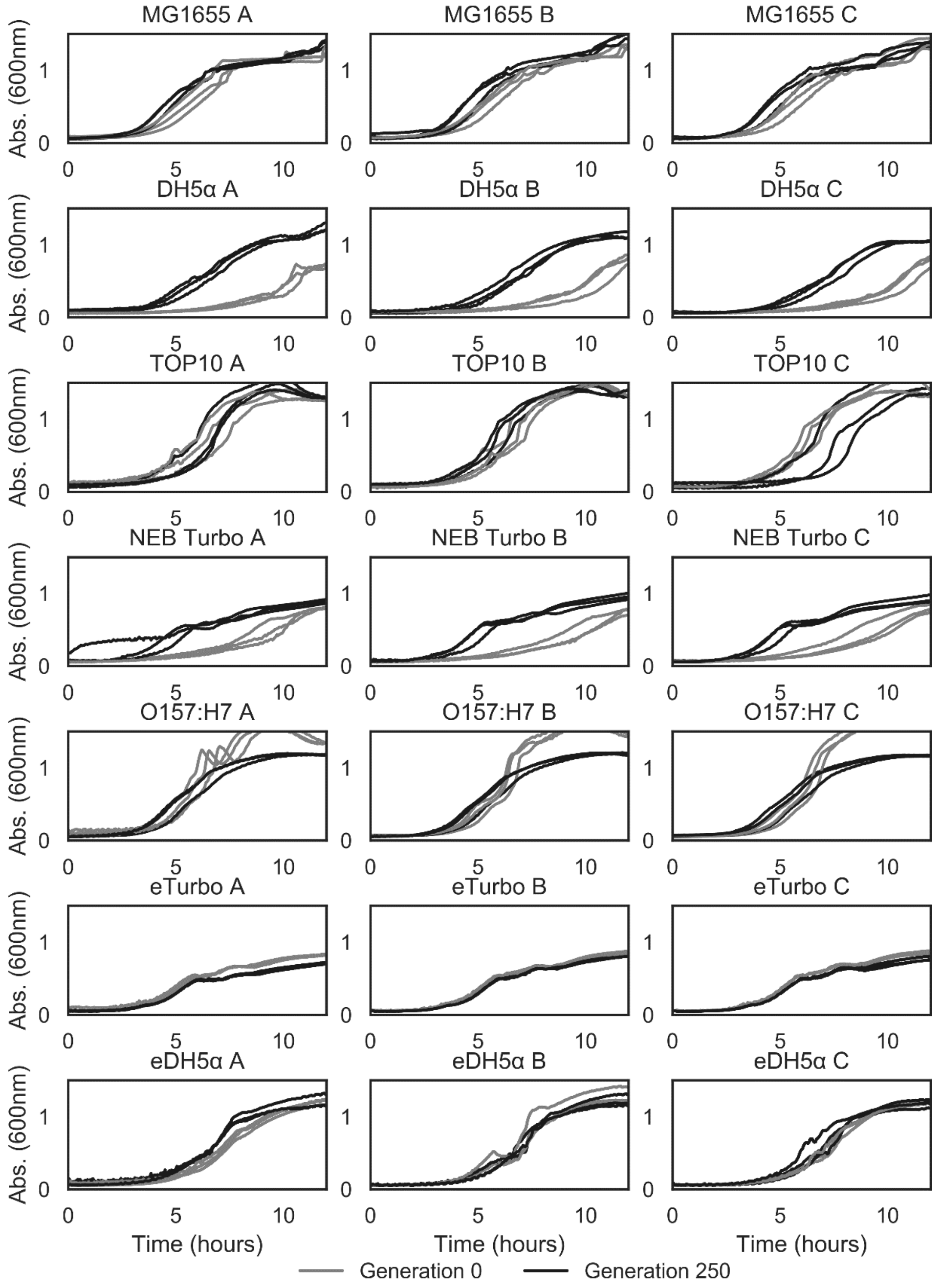

**Supplementary Figure 3**: Growth curves of *E. coli* strains at generation 0 and generation 250 of competitive fitness assays. *E. coli* strains containing pRPU and grown in M9 media with kanamycin. Growth curves were measured from samples of each culture at generation 0 (gray curves) and generation 250 (black curves). Each biological replicate (A, B, C) was measured in technical triplicate. Some curves overlap.

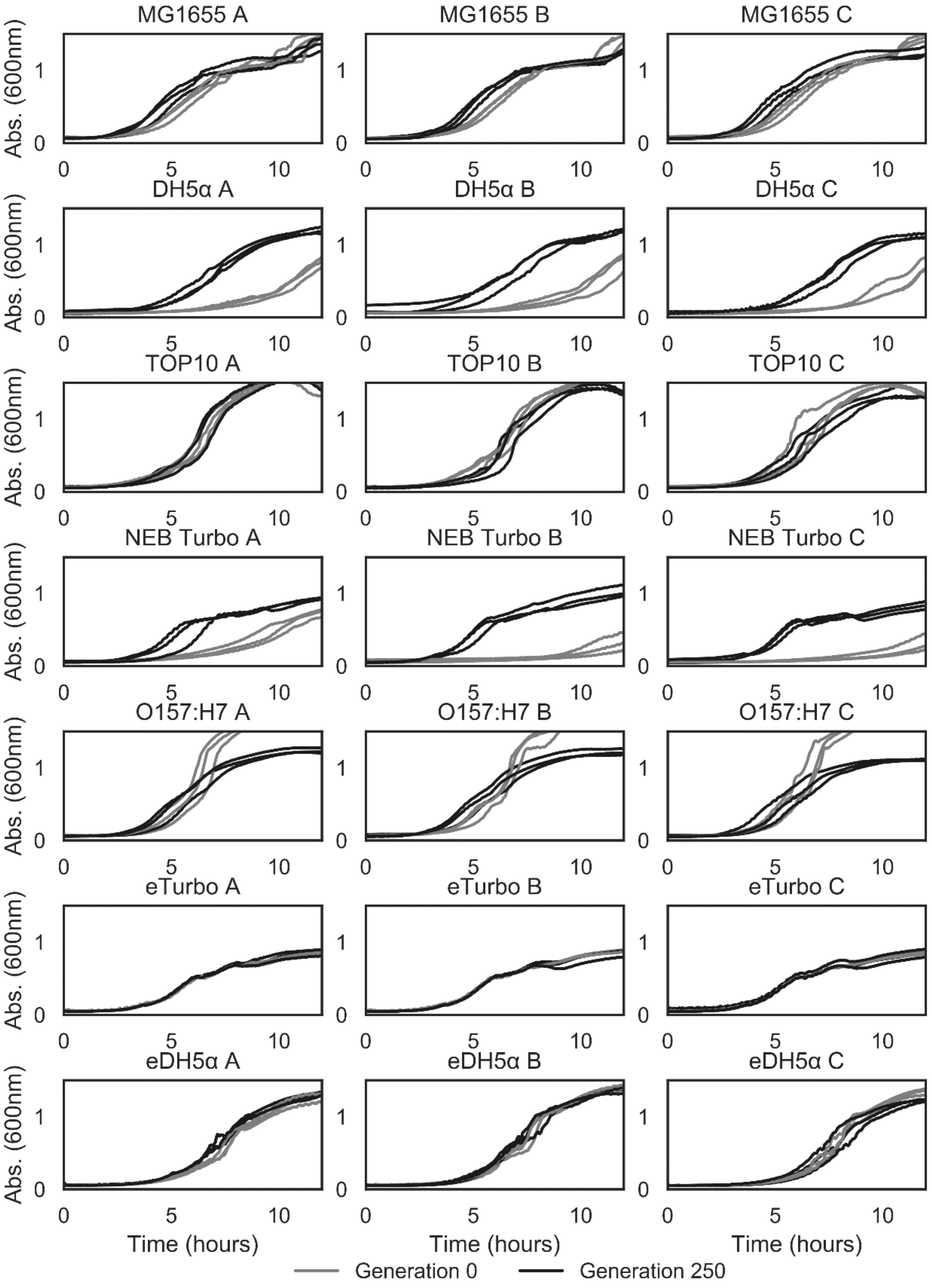

**Supplementary Figure 4**: Growth curves show increased growth rates after adapting *E. coli* strains to M9 media. Growth curves of NEB Turbo (left column) and DH5α (right column) before adaptation (gray curves) and after 100 generations of adaptation (black curves). Growth curves were measured in LB media (upper plots) and M9 media (lower plots). Measured in biological triplicate.

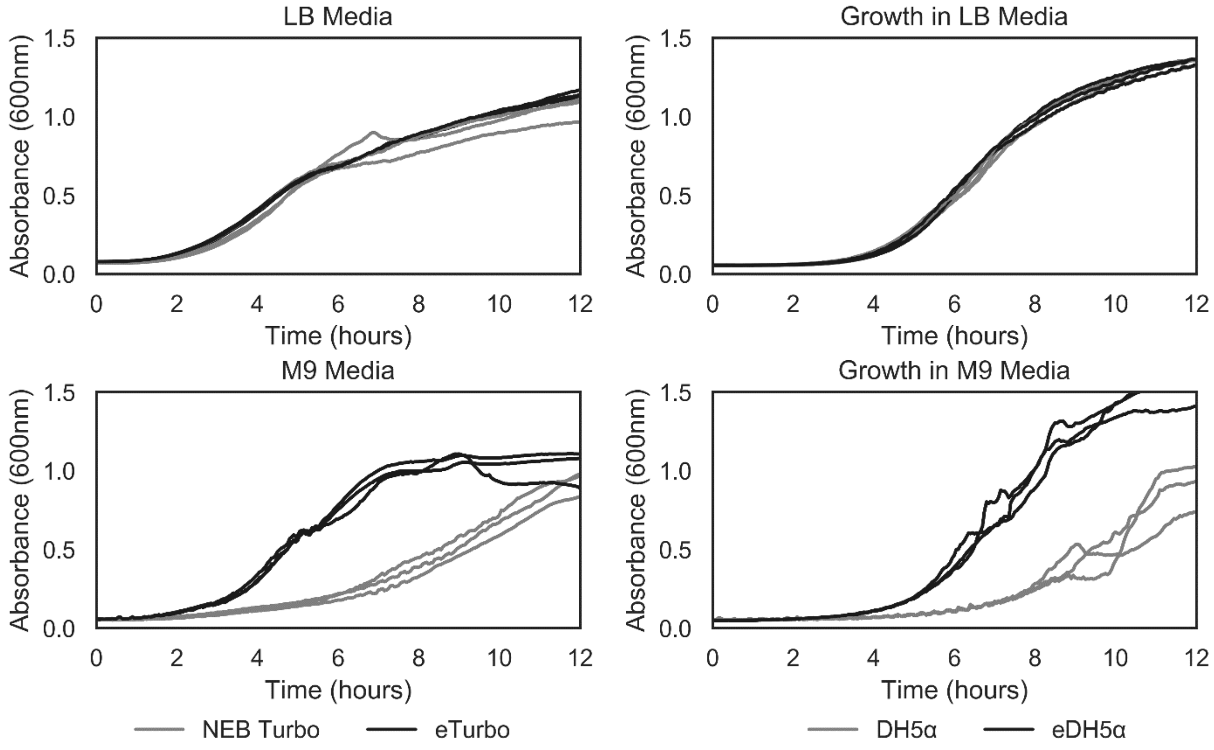

**Supplementary Figure 5**: Automated gating to isolate single cell event in flow cytometry. Cytometry data was gated by automating the subtraction of background events. Left, control blank sample to identify background events. Center, raw data from cytometer containing background and cell events. Right, gated results show cell events with background events removed.

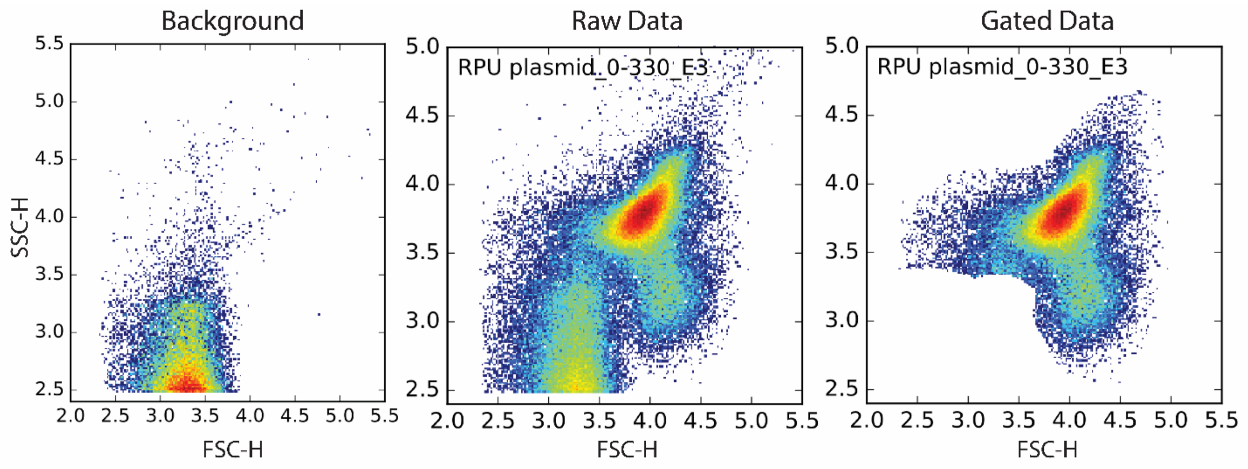

**Improved stability of an engineered function using adapted bacterial strains**

**Supplementary table 1**: Variability (geometric standard deviation, $\sigma_g$) of YFP expression from pRPU in different strains of *E. coli*.

| Strain | Variability ($\sigma_g$) |
|---|---|
| MG1655 | 1.14 |
| O157:H7 | 1.20 |
| TOP10 | 1.06 |
| DH5α | 1.34 |
| eDH5α | 1.16 |
| NEB Turbo | 1.12 |
| eTurbo | 1.08 |

**Supplementary table 2**: Growth rate (as doublings per hour) with standard deviation of seven *E. coli* strains at generation 0 ($\mu_0$) and generation 250 ($\mu_{250}$) of the stability assays and competitive fitness assays.

| | Stability Assay | | | | | | | |
|---|---|---|---|---|---|---|---|---|
| | $\mu_0$ | $\mu_{250}$ | $N_0$ | $N_{250}$ | $\mu_0$ | $\mu_{250}$ | $N_0$ | $N_{250}$ |
| MG1655 | 1.36±0.08 | 1.65±0.11 | 9 | 8 | 1.26±0.14 | 1.52±0.17 | 8 | 9 |
| O157:H7 | 1.47±0.12 | 1.60±0.06 | 9 | 9 | 1.52±0.07 | 1.60±0.06 | 9 | 9 |
| TOP10 | 1.15±0.22 | 1.27±0.24 | 9 | 9 | 1.25±0.14 | 1.25±0.15 | 9 | 9 |
| DH5α | 0.71±0.09 | 1.26±0.12 | 9 | 9 | 0.77±0.08 | 1.15±0.15 | 9 | 9 |
| eDH5 | 1.16±0.23 | 1.12±0.16 | 9 | 8 | 1.17±0.03 | 1.36±0.10 | 9 | 9 |
| NEB Turbo | 0.65±0.06 | 1.26±0.08 | 9 | 8 | 0.63±0.07 | 1.25±0.08 | 9 | 9 |
| eTurbo | 1.12±0.10 | 1.13±0.05 | 9 | 9 | 1.04±0.08 | 1.14±0.04 | 9 | 9 |

**Supplementary table 3**: Growth rate (as doublings per hour) with standard deviation of NEB Turbo and eTurbo in LB and M9 growth media. The change in growth rate (Δμ) calculated using $\mu_{eTurbo}/\mu_{NEB\ Turbo}$.

| | Growth Rate ($hr^{-1}$) | | |
|---|---|---|---|
| Media | NEB Turbo | eTurbo | Δμ |
| LB | 1.87±0.19 | 1.92±0.23 | 1.03±0.16 |
| M9 | 0.62±0.08 | 1.01±0.16 | 1.63±0.34 |

**Supplementary Table 4**: Growth rate (as doublings per hour) with standard deviation of DH5α and eDH5α in LB and M9 growth media. The change in growth rate (Δμ) calculated using $\mu_{eDH5\alpha}/\mu_{DH5\alpha}$.

| | Growth Rate ($hr^{-1}$) | | |
|---|---|---|---|
| Media | DH5α | eDH5α | Δμ |
| LB | 1.63±0.07 | 1.54±.04 | 0.94±0.05 |
| M9 | 0.70±0.11 | 1.23±.10 | 1.75±0.31 |

**Supplementary Table 5**: Minimum Information Standard for Engineered Organism Experiments (MIEO).

| **MIEO Category** | **Factor** | **Level** | **Supplier Part** |
|---|---|---|---|
| Media components (M9) | KH2PO4 | 3 g/L | BD (248510) |
| | Na2HPO4 | 6.78 g/L | BD (248510) |
| | NaCl | 0.5 g/L | BD (248510) |
| | NH4Cl | 1.0 g/L | BD (248510) |
| | D-glucose | 0.721 g/L | Sigma (G8270) |
| | Casamino acids | 2 g/L | Calbiochem(2240) |
| | CaCl2 | 0.011 g/L | Sigma (21115) |
| | MgSO4 | 0.241 g/L | Sigma (83266) |
| | Vitamin B1 (Thiamine) | 3.40 x 10-4 g/L | Sigma-Aldrich (T4625) |
| Media properties | pH | 7.4 | |
| | LB-Miller | 25g/L | Fisher BioReagents (BP1426) |
| | Bacto-agar | 15g/L` | BD (214010) |
| Container geometry | Type | "Culture" tube | Falcon (352059) |
| | Container shape | Round | |
| | Container bottom | Round | |
| | Container volume | 14 mL | |
| | Fill volume | 3mL | |
| | Cover | Snap cap | |
| Container shaking | Shaking speed | 200 rpm | |
| | Shaking diameter | 2.5 cm | |
| | Shaking mode | Orbital | |
| Time | Growth time | 24h | |
| Environment | Temperature | 37°C | |
| | Relative humidity | Not measured | |
| Selective Agents | Antibiotic type | Kanamycin | gibco (11815-32) |
| | Antibiotic concentration | 50 ug/mL | |
| Inoculum | Type | Single colony | |
| | Concentration at inoculation | N/A | |
| | Age of inoculum at inoculation | N/A | |

| | | | |
|---|---|---|---|
| Passage | Volume of inoculum | 3μL | |
| | Age of inoculum at inoculation | 24 hours | |
| | Concentration at inoculation | 10^-3 | |
| Flow Cytometry | Device | Cytometer | |
| | Device | Autosampler | |
| | PBS | | Invitrogen (AM9625) |
| | Focusing Fluid | | Attune Focusing Fluid (4488621) |
| | Calibration Beads | | Spherotech (RCP-30-5A) |
| | Chloramphenicol | | Sigma(C1919) |

**pRPU Plasmid Sequence**:

L3S3P21 – red text
$P_{J23101}$ – green highlighted text
RiboJ – gray highlighted text
**RBS – green emboldened text**
eYFP – yellow highlighted text
$T_{L3S2P21}$ – gray highlighted red text
P15A – cyan highlighted text
Kanamycin resistance – underlined text

>plasmid pRPU
CCAATTATTGAAGGCCTCCCTAACGGGGGGCCTTTTTTTGTTTCTGGTCTCCCGCTTGATAAGT
CCCTAACTTTTACAGCTAGCTCAGTCCTAGGTATTATGCTAGCCTGAAGCTGTCACCGGATGTG
CTTTCCGGTCTGATGAGTCCGTGAGGACGAAACAGCCTCTACAAATAATTTTGTTTAATACTAG
AG**AAAGAGGGGAAA**TACTAGATGGTGAGCAAGGGCGAGGAGCTGTTCACCGGGGTGGTGCCCAT
CCTGGTCGAGCTGGACGGCGACGTAAACGGCCACAAGTTCAGCGTGTCCGGCGAGGGCGAGGGC
GATGCCACCTACGGCAAGCTGACCCTGAAGTTCATCTGCACCACAGGCAAGCTGCCCGTGCCCT
GGCCCACCCTCGTGACCACCTTCGGCTACGGCCTGCAATGCTTCGCCCGCTACCCCGACCACAT
GAAGCTGCACGACTTCTTCAAGTCCGCCATGCCCGAAGGCTACGTCCAGGAGCGCACCATCTTC
TTCAAGGACGACGGCAACTACAAGACCCGCGCCGAGGTGAAGTTCGAGGGCGACACCCTGGTGA
ACCGCATCGAGCTGAAGGGCATCGACTTCAAGGAGGACGGCAACATCCTGGGGCACAAGCTGGA
GTACAACTACAACAGCCACAACGTCTATATCATGGCCGACAAGCAGAAGAACGGCATCAAGGTG
AACTTCAAGATCCGCCACAACATCGAGGACGGCAGCGTGCAGCTCGCCGACCACTACCAGCAGA
ACACCCCAATCGGCGACGGCCCCGTGCTGCTGCCCGACAACCACTACCTTAGCTACCAGTCCGC
CCTGAGCAAAGACCCCAACGAGAAGCGCGATCACATGGTCCTGCTGGAGTTCGTGACCGCCGCC
GGGATCACTCTCGGCATGGACGAGCTGTACAAGTAACTCGGTACCAAATTCCAGAAAAGAGGCC
TCCCGAAAGGGGGGCCTTTTTTCGTTTTGGTCCGATCCTCTACGCCGGACGCATCGTGGCCGGC
ATCACCGGCGCCACAGGTGCGGTTGCTGGCGCCTATATCGCCGACATCACCGATGGGGAAGATC

GGGCTCGCCACTTCGGGCTCATGAGCAAATATTTTATCTGAGGTGCTTCCTCGCTCACTGACTC
GCTGCACGAGGCAGACCTCAGCGCTAGCGGAGTGTATACTGGCTTACTATGTTGGCACTGATGA
GGGTGTCAGTGAAGTGCTTCATGTGGCAGGAGAAAAAAGGCTGCACCGGTGCGTCAGCAGAATA
TGTGATACAGGATATATTCCGCTTCCTCGCTCACTGACTCGCTACGCTCGGTCGTTCGACTGCG
GCGAGCGGAAATGGCTTACGAACGGGGCGGAGATTTCCTGGAAGATGCCAGGAAGATACTTAAC
AGGGAAGTGAGAGGGCCGCGGCAAAGCCGTTTTTCCATAGGCTCCGCCCCCCTGACAAGCATCA
CGAAATCTGACGCTCAAATCAGTGGTGGCGAAACCCGACAGGACTATAAAGATACCAGGCGTTT
CCCCCTGGCGGCTCCCTCGTGCGCTCTCCTGTTCCTGCCTTTCGGTTTACCGGTGTCATTCCGC
TGTTATGGCCGCGTTTGTCTCATTCCACGCCTGACACTCAGTTCCGGGTAGGCAGTTCGCTCCA
AGCTGGACTGTATGCACGAACCCCCCGTTCAGTCCGACCGCTGCGCCTTATCCGGTAACTATCG
TCTTGAGTCCAACCCGGAAAGACATGCAAAAGCACCACTGGCAGCAGCCACTGGTAATTGATTT
AGAGGAGTTAGTCTTGAAGTCATGCGCCGGTTAAGGCTAAACTGAAAGGACAAGTTTTGGTGAC
TGCGCTCCTCCAAGCCAGTTACCTCGGTTCAAAGAGTTGGTAGCTCAGAGAACCTTCGAAAAAC
CGCCCTGCAAGGCGGTTTTTTCGTTTTCAGAGCAAGAGATTACGCGCAGACCAAAACGATCTCA
AGAAGATCATCTTATTAAGGGGTCTGACGCTCAGTGGAACGAAAAATCAATCTAAAGTATATAT
GAGTAAACTTGGTCTGACAGTTACCTTAGAAAAACTCATCGAGCATCAAATGAAACTGCAATTT
ATTCATATCAGGATTATCAATACCATATTTTTGAAAAAGCCGTTTCTGTAATGAAGGAGAAAAC
TCACCGAGGCAGTTCCATAGGATGGCAAGATCCTGGTATCGGTCTGCGATTCCGACTCGTCCAA
CATCAATACAACCTATTAATTTCCCCTCGTCAAAAATAAGGTTATCAAGTGAGAAATCACCATG
AGTGACGACTGAATCCGGTGAGAATGGCAAAAGCTTATGCATTTCTTTCCAGACTTGTTCAACA
GGCCAGCCATTACGCTCGTCATCAAAATCACTCGCATCAACCAAACCGTTATTCATTCGTGATT
GCGCCTGAGCGAGACGAAATACGCGATCGCTGTTAAAAGGACAATTACAAACAGGAATCGAATG
CAACCGGCGCAGGAACACTGCCAGCGCATCAACAATATTTTCACCTGAATCAGGATATTCTTCT
AATACCTGGAATGCTGTTTTCCCGGGGATCGCAGTGGTGAGTAACCATGCATCATCAGGAGTAC
GGATAAAATGCTTGATGGTCGGAAGAGGCATAAATTCCGTCAGCCAGTTTAGTCTGACCATCTC
ATCTGTAACATCATTGGCAACGCTACCTTTGCCATGTTTCAGAAACAACTCTGGCGCATCGGGC
TTCCCATACAATCGATAGATTGTCGCACCTGATTGCCCGACATTATCGCGAGCCCATTTATACC
CATATAAATCAGCATCCATGTTGGAATTTAATCGCGGCCTGGAGCAAGACGTTTCCCGTTGAAT
ATGGCTCATAACACCCCTTGTATTACTGTTTATGTAAGCAGACAGTTTTATTGTTCATGATGAT
ATATTTTTATCTTGTGCAATGTACATCAGAGATTTTGAGACACAA
